# A novel Mn-dependent peroxidase contributes to tardigrade anhydrobiosis

**DOI:** 10.1101/2020.11.06.370643

**Authors:** Yuki Yoshida, Tadashi Satoh, Chise Ota, Sae Tanaka, Daiki D. Horikawa, Masaru Tomita, Koichi Kato, Kazuharu Arakawa

## Abstract

Tardigrades are microscopic animals that are capable of tolerating extreme environments by entering a desiccated ametabolic state known as anhydrobiosis. While antioxidative stress genes, antiapoptotic pathways and tardigrade-specific intrinsically disordered proteins have been implicated in the anhydrobiotic machinery, conservation of these mechanisms is not universal within the phylum Tardigrada, suggesting the existence of overlooked components. Here, we show that a novel Mn-dependent peroxidase is an important factor in tardigrade anhydrobiosis. Through comparative time-series transcriptome analysis of *Ramazzottius varieornatus* specimens exposed to desiccation or ultraviolet light, we first identified several novel gene families without similarity to existing sequences that are induced rapidly after stress exposure. Among these, a single gene family with multiple orthologs that is highly conserved within the phylum Tardigrada and enhances oxidative stress tolerance when expressed in human cells was identified. Crystallographic study of this protein suggested Zn or Mn binding at the active site, and we further confirmed that this protein has Mn-dependent peroxidase activity *in vitro*. Our results demonstrated novel mechanisms for coping with oxidative stress that may be a fundamental mechanism of anhydrobiosis in tardigrades. Furthermore, localization of these sets of proteins in the Golgi apparatus suggests an indispensable role of the Golgi stress response in desiccation tolerance.

## Introduction

Tardigrades are microscopic animals that are renowned for their ability to enter an ametabolic state known as cryptobiosis [1] or, more particularly, anhydrobiosis (life without water), which is cryptobiosis upon almost complete desiccation. Tardigrades can withstand extreme conditions in this dormant state, including extreme temperature, pressure, high doses of ionizing radiation, and exposure to the vacuum of space [2-8], yet they quickly resume life upon rehydration. Anhydrobiosis has been acquired in multiple lineages encompassing all kingdoms of life, but tardigrades are unique in multi-cellular animals that they can enter anhydrobiosis within minutes[9], and that the mechanism does not rely on trehalose and LEA proteins [10, 11]. The molecular machinery of tardigrade anhydrobiosis is beginning to be uncovered due to the availability of genomic resources [12-14], leading to the identification of several tardigrade-specific genes, such as CAHS, SAHS, MAHS, LEAM, and Dsup, that have been suggested to play critical roles in cellular protection upon anhydrobiosis [14-16]. Notably, Dsup is a nucleus-localizing DNA-binding protein that is reported to protect DNA molecules from hydroxyl radicals, where the induction of this single protein in mammalian cells and plants can increase its radiation tolerance [14, 17-19]. However, these proteins are not conserved across the phylum Tardigrada (as they are not conserved in the class Heterotardigrada) [20], and the necessary and sufficient set of genes and pathways enabling anhydrobiosis remains elusive.

Of the many adverse extremities tardigrades can tolerate in anhydrobiosis, radiation is unique in that tardigrades can better tolerate it in the active hydrated state than in the inactive desiccated state[21, 22], suggesting the existence of efficient repair pathways in addition to the protective mechanisms identified thus far. Tolerance to radiation in tardigrades is a cross-tolerance of anhydrobiosis [23], and the overlapping pathway is presumably the defense against reactive oxygen species (ROS) that mediates protein oxidation and DNA damage [24, 25]. To this end, we employed ultraviolet C (UVC), a low-level energy stressor that causes oxidative stress, to screen for tardigrade-unique components for ROS defense in *Ramazzottius varieornatus*. Tardigrades are capable of tolerating approximately 1,000-fold higher dosages of ultraviolet B (UVB) and UVC than human cell lines [6, 26].

## Results

### Transcriptome sequencing of UVC exposed *R. varieornatus*

Previous studies have observed that *R. varieornatus* exposed to 2.5 kJ/m^2^ UVC showed a prolonged decrease in movement for approximately 1 day, which presumably is the critical period for the ROS response. We first validated this observation on a finer time scale, where individuals who were exposed to 2.5 kJ/m^2^ showed significantly lower movements from 2-9 hours after exposure (**Fig 1A**).We then conducted transcriptome profiling from 0-12 hours and 0-72 hours to screen for genes induced in this period (**S1 Table**). Initial clustering of expression values using Spearman correlation indicated that the transcriptome profiles drastically shifted between 3-4- and 24-36-hours post exposure (**S2 Fig**), suggesting consistency with the motility of the animals. We found a total of 3,324 differentially expressed genes following exposure to UVC (DESeq2, FDR<0.05), of which 1,314 and 2,110 genes to be up-regulated and down-regulated, accordingly (**S2 Fig**). Genes with high fold change (>4) were comprised of various genes, including chaperones (Mitochondrial chaperone BCS1), DNA damage repair pathways (XRCC4, PARP), metalloproteases (NAS-13), anti-oxidative pathways (GST), and previously identified tardigrade specific protection-related genes (CAHS, SAHS). Interestingly, DEGs that had high expression values included several of those with high fold change, additionally anti-oxidative stress genes thioredoxin and Peroxiredoxin (**S3 Table, S4 Table**). These genes were induced during tardigrade anhydrobiosis [13], suggesting similar pathways are being regulated between desiccation and UVC exposure. Additionally, we found that the zinc metalloprotease NAS, apolipoproteins, autophagy-related sequestosome, and the mitochondrial chaperone BCS1 were highly expressed (TPM>1000) after exposure (**S5 Text**).

**Fig 1.**
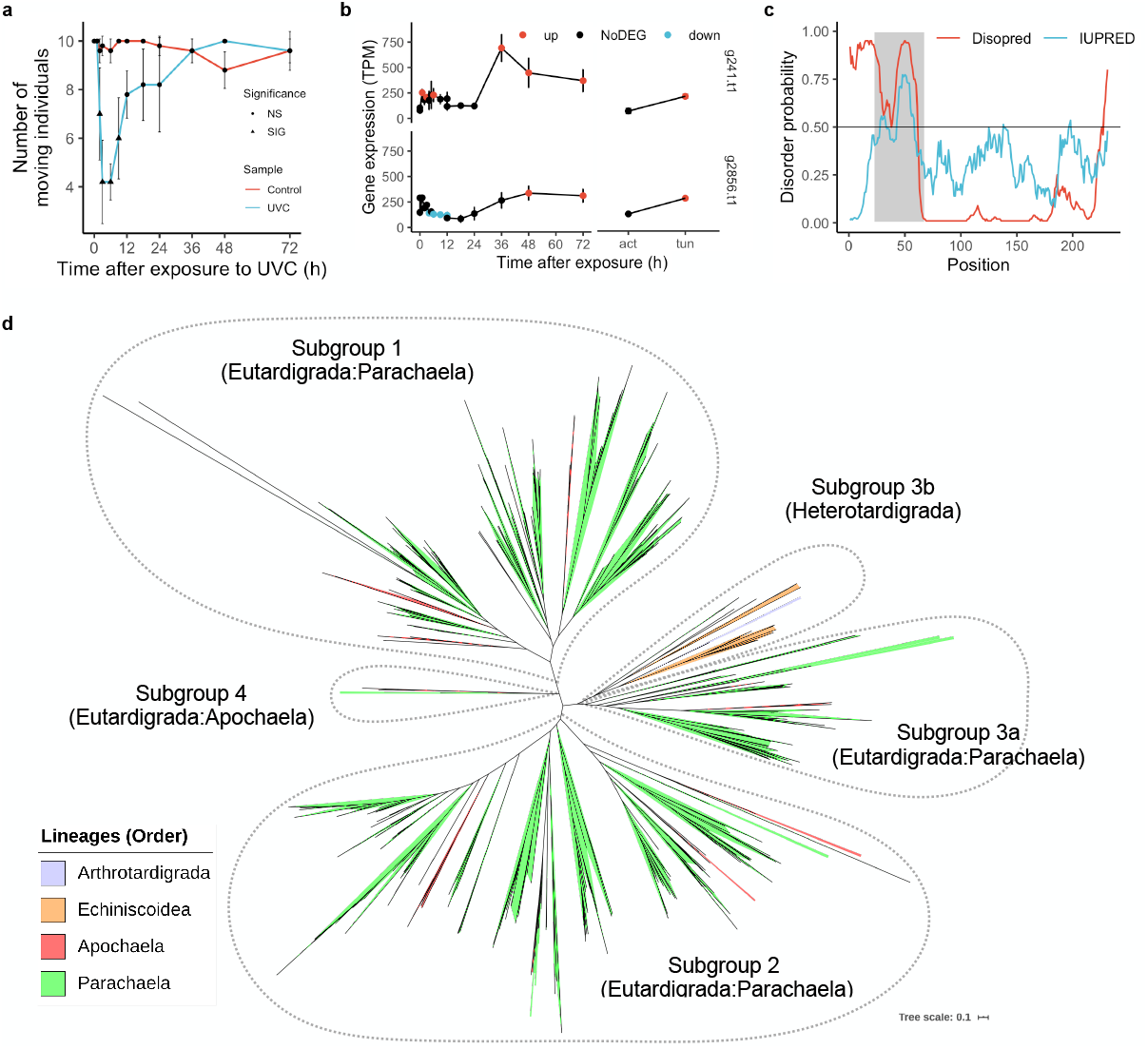
Identification of a novel stress responsive gene family conserved within Tardigrada phylum. [a] *R. varieornatus* specimens were exposed to 2.5kJ/m^2^ of UVC and movements were observed at each time point. Asterisks indicate significant differences (Tukey HSD test, p-value < 0.05). [b] Expression values of two tardigrade-specific genes with no known annotations. [c] Probability plot of disordered regions in g12777 predicted by DISOPRED or IUPRED. Disordered regions predicted are indicated with a gray highlight. [d] Phylogenetic tree of g12777 orthologs in Tardigrada. Each ortholog was colored by the corresponding linage. Four subfamilies were found and named according to the major lineage.

### Identification of the g12777 gene family as a novel stress responsive gene family

To screen for genes responsible for the cross-tolerance of anhydrobiosis and UVC exposure, we analyzed the intersection of DEGs in the above result and our previous differential transcriptome analysis during slow desiccation. We found 141 genes that were upregulated in both conditions (**S7 Table** that were significantly enriched in Gene Ontology terms related to antioxidative stress (*e*.*g*., glutathione transferase activity, superoxide dismutase activity, etc., **S8 Table**), and these genes also included the previously identified tardigrade-specific heat soluble proteins CAHS and SAHS. Seventy-five of these genes were hypothetical genes, and only two (g2856/RvY_14843, g241/RvY_00334) contained no known functional domains (Interproscan, CDD, Pfam-A, Superfamily, SMART, **Fig 1b, S9 Text, S10 Table, S11 Table**). The g241 gene showed similarity with bacterial genes, and phylogenetic analysis of this gene suggested a possible horizontal gene transfer event at the early stages of the Tardigrada lineage (**S16 Fig, S17 Text**). The remaining g2856 was strongly multiplied within the *R. varieornatus* genome (total of 35 copies), and several of the orthologs were duplicated in tandem (**S21 Table**) and were highly conserved throughout the Tardigrada phylum, including in nonanhydrobiotic Heterotardigrada species. Phylogenetic analysis of g2856 orthologs indicated that there are 4 subgroups (**Fig 1c**), where a single subfamily was comprised of Heterotardigrada species. Within the 35 copies of the g2856 gene family, the g12777 gene was found to have 2.5-fold induction of expression during slow desiccation of *R. varieornatus*. Additionally, informatics-based analysis predicted a signal peptide and a disordered region in the N-terminus of this protein (**Fig 1d**), similar to various tardigrade-specific anhydrobiosis-related genes. These characteristics suggested that this gene may play an important role in tardigrade anhydrobiosis; therefore, we submitted this g12777 gene for further functional analysis.

### g12777 has a highly conserved Mn^2+^ biding site

We crystallized the putative globular domain that lacked the N-terminus 62 amino acids of g12777 protein and solved the crystal structures as two forms containing Mn^2+^ or Zn^2+^ ions at 2.30 Å and 1.60 Å resolutions, respectively (**Fig 2a, S22abcd Fig, S23 Table, S24 Text**). These crystallographic data revealed that g12777 protein possesses a β-sandwich domain sharing a common Mn^2+^ and Zn^2+^ binding site. In this site, a Mn^2+^ ion was coordinated by three aspartic acids with high electrostatic surface potential (**Fig 2bc**). In the Mn^2+^-binding site located at the negatively charged patch comprising β1–β2 and β3–β4 loops, the side-chain oxygen atoms of Asp92, Asp98, and Asp131 and the main-chain carbonyl atom of Ala96 and two water molecules coordinate with Mn^2+^ ion at distances of 2.2–2.9 Å (**Fig 2c**). As for the Zn^2+^-binding site located at the almost same region as that of Mn^2+^, the side-chain oxygen atoms of Asp92, Asp98, Asp161, and Asp163 coordinate with Zn^2+^ ion at distances of 1.9–2.0 Å (**S22d Fig**). Considerable conformational difference was observed between Mn^2+^- and Zn^2+^-bound forms in terms of the metal-binding site, in which Asp92 and Asp98 were commonly involved in Mn^2+^ and Zn^2+^ binding (**Fig 2c** and **S22d Fig**). These aspartic acid residues were highly conserved within tardigrade orthologs (**S22ef Fig**), and the highly conserved CD-CD motif containing two Asp residues (D92 and D98, **S22efg F**ig) was used in the metal ion binding in both structures, suggesting the importance of these residues. The cysteine residues in this motif formed a disulfide bond in the crystal structure, suggesting that this disulfide bond may also contribute to this protein’s function. We also observed another region that was highly conserved (W194 and R216, **S22ef Fig**), suggesting that another functional site may be present.

**Fig 2.**
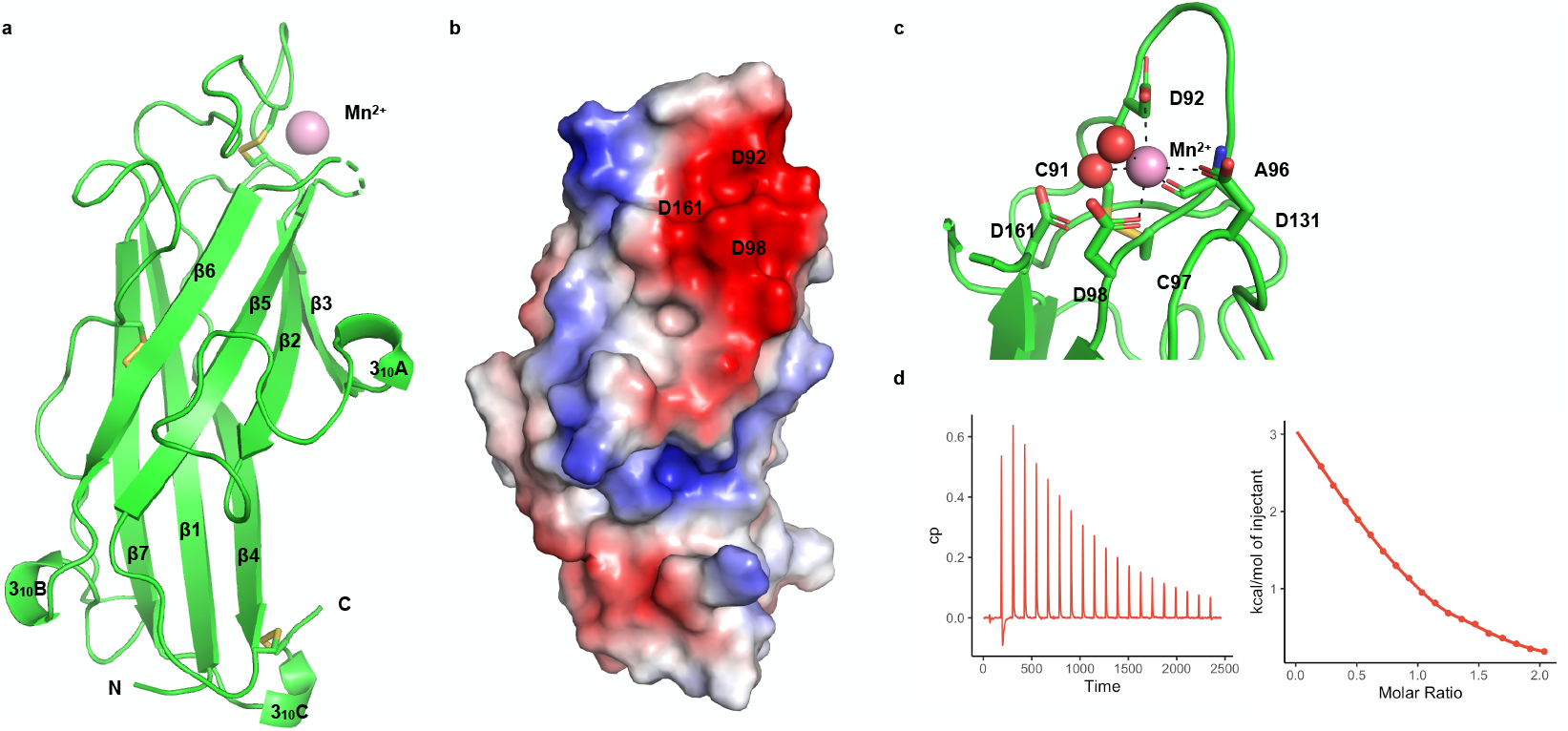
g12777 protein has a SOD-like-sandwich fold and binds to Mn^2+^ ion. [a] The crystal structure of the catalytic domain of the g12777 protein. Bound Mn^2+^ ion and residues involved in disulfide bond formation are indicated with a pink sphere and stick, respectively. Positions of the N and C termini are indicated as letters. [b] Electrostatic surface potential of g12777 protein. The surface model of g12777 is colored according to the electrostatic surface potential (blue, positive; red, negative; scale from -50 to +50 kT/e). [c] Close-up view of the Mn^2+^-binding site. The binding site is comprised from three aspartic acid residues (D92, D98, and D131) and a disulfide bond with close proximity (C91 and C97). [d] Mn^2+^ binding affinity measured by isothermal titration calorimetry. The left panel indicates the raw data, while the right panel represents the integrated heat values corrected for the heat of dilution and fit to a one-site binding model (solid line).

ITC experiment demonstrated that the estimated dissociation constants of catalytic domain of g12777 and Zn^2+^, Mn^2+^, and Ca^2+^ were 1.92×10^−6^±6.00×10^−8^ M, 2.42×10^−5^±1.75×10^−6^ M, or 1.39×10^−4^±3.02×10^−5^ M, respectively, indicating that the binding affinity of Zn^2+^ was the highest among these divalent cations (**Fig 2d, S22g Fig, S25 Table**). Importantly, the binding stoichiometry (*n*) of Zn^2+^, Mn^2+^, and Ca^2+^ were 1.00±0.00, 0.77±0.02, and 0.80±0.17, indicating that catalytic domain of g12777 binds to the one metal ions in solution (**Fig 2d, S22g Fig, S25 Table)**.

### g12777 is a Mn^2+^ dependent peroxidase

To validate where this protein functions in cultured cells, we constructed a GFP-tagged recombinant g12777 protein in the pAcGFP1-N1 plasmid and expressed these proteins in the HEK293 cell line. Costaining with DAPI and CellLight GolgiRFP suggested that this protein localizes in the Golgi apparatus (**Fig 3a**). Six of the most highly expressed orthologs of the g12777 family of proteins (highly expressed in the UVC time course and differentially expressed during slow-dry anhydrobiosis) also showed Golgi localization (**S26 Fig**). To validate this protein’s antioxidative stress functions, we subjected cells expressing GFP-tagged g12777 proteins to hydrogen peroxide (H_2_O_2_) exposure. The transfected cells showed increased survival at 0.2-0.3 mM H_2_O_2_ as measured by MTT assays (**Fig 3b**). Similar results were obtained (0.1-0.2 mM H_2_O_2_) by flow cytometry measurements (Annexin V-negative + SYTOX Blue-negative cells, **Fig 3c**). Substitution of the Asp residues comprising Zn-1 to alanine (AMNP-D2A) showed a decrease in cell survival (**Fig 3bc**), indicating that this Mn-1 binding site is required for antioxidative stress function. We then assayed whether recombinant proteins also have an antioxidative stress function. Recombinant β-sandwich domain of g12777 protein showed significant peroxidase activity when Mn^2+^ ions but not Zn^2+^ ions were present (**Fig 3d**). The peroxidase activity of AMNP in Mn2+ conditions were 1/20 of that of bovine liver catalase (**S27 Fig**). Removal of the signal peptide and disorder region did not significantly reduce tolerance against oxidative stress (**Fig 3bc**), suggesting that the disordered region does not directly contribute to the anti-oxidative stress function.

**Fig 3.**
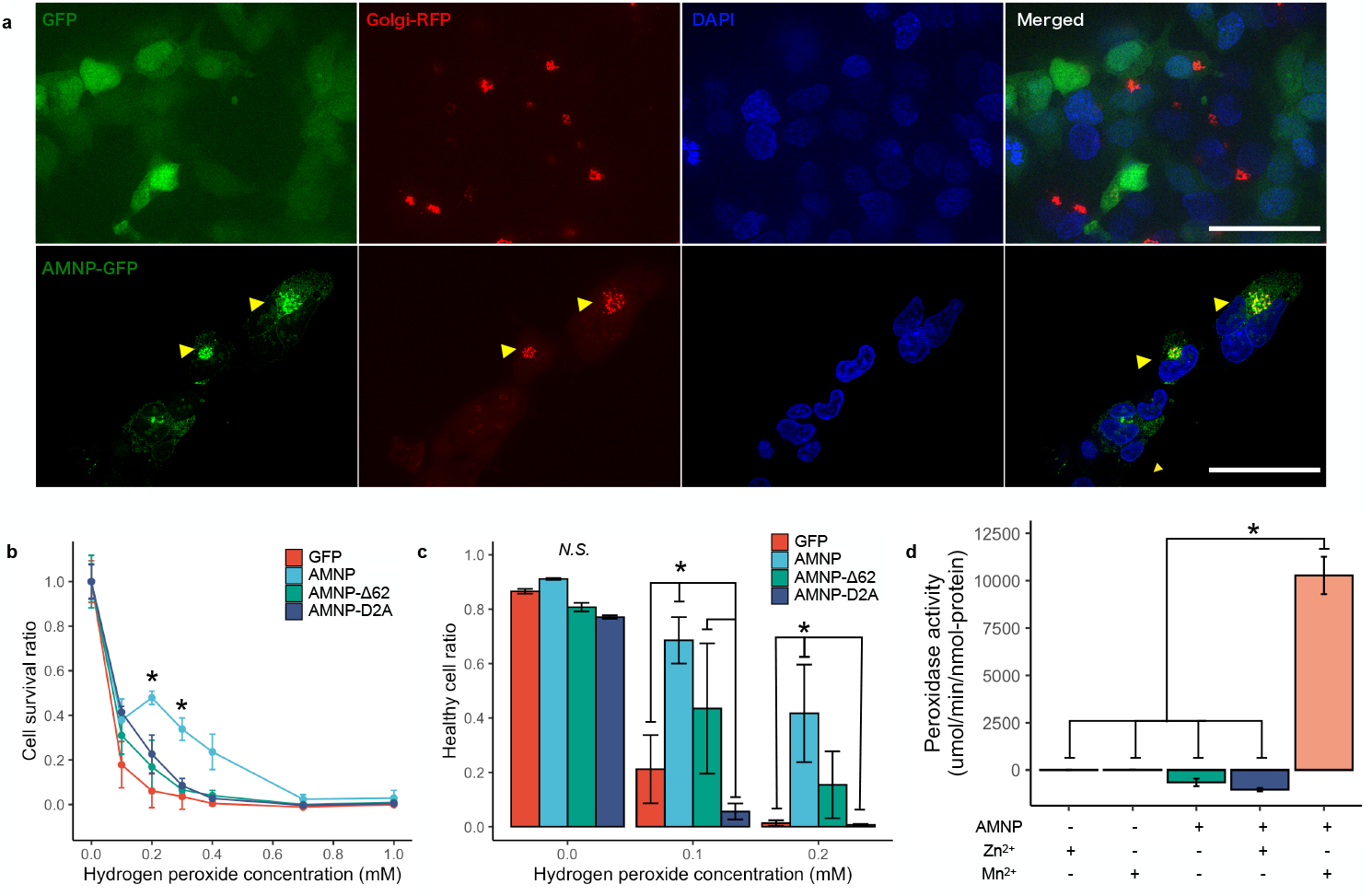
AMNP is a Golgi apparatus localizing protein that enhances anti-oxidative stress. [a] The g12777 protein was fused to GFP and transfected to HEK293 cells. Cells were co-transfected with CellLight Golgi-RFP and DAPI. Yellow arrows indicate the co-localization of g12777 and Golgi-RFP. Scale bar 25µm. [b] Cell transfected with g12777-GFP were exposed to hydrogen peroxide for 30 minutes and subjected to MTT assay after 24 hours incubation. Only the full-length g12777-GFP show an increase in survival at around 0.2-0.3 mM. ANOVA + Tukey HSD (FDR < 0.05). [c] Cells transfected with g12777-GFP were exposed to hydrogen peroxide for 30 minutes and subjected to MACS flow cytometry to detect SYTOX blue and AnnexinV-Alexa 657 fluorescence after 24 hours incubation. Only full-length g12777-GFP shows an increase in survival. ANOVA + Tukey HSD (FDR <0.05). [d] Peroxidase activity of the recombinant protein lacking the N terminal 62aa region. Peroxidase function was present only when manganese ions were present. Error bars indicate standard errors.

## Discussion

These findings suggest that this protein is a manganese-dependent peroxidase, independent of other components of the cell. Hence, we named this gene family Anhydrobiosis-related Mn-dependent Peroxidase (AMNP). We hypothesize that the presence of Mn-1 and the disulfide bond at the active site is suggestive of the utilization of both mechanisms for peroxidase activity, which could categorize AMNP as a new class of peroxidase. Constructs lacking the disordered region showed anti-oxidative stress functions. Intrinsically disordered proteins (IDP) are implied to contribute to cellular tolerance in Tardigrades [15, 16] and Drosophila [27]. IDPs have been proposed to participate in protein stability [28], therefore we hypothesize this disordered region may contribute to protein stability of AMNP proteins. We have also observed that the AMNP gene utilizes Mn^2+^ ions for peroxidase function, while it has a higher affinity for Zn^2+^ ions. We speculate that this affinity is related to the localization of this protein in the Golgi apparatus, since the Golgi apparatus has a strictly controlled high manganese concentration [29-32], suggesting the antioxidative stress functionality in the Golgi seems to be crucial for anhydrobiotic survival. The importance of the Golgi apparatus in stress response and regulation has been studied [33-35], but its contribution to anhydrobiosis has not been discussed in detail to date. We believe that the results of this work can be a starting point for a more comprehensive study of the intracellular mechanisms of anhydrobiosis.

In conclusion, our results provide a global image of the transcriptomic response against UVC in an extremophile tardigrade and showed that the oxidative stress response partly comprised of the novel Mn-dependent peroxidase family AMNP is one of the central mechanisms in tardigrade anhydrobiosis. Moreover, AMNP is the first tardigrade-specific anhydrobiosis-related gene that is conserved throughout the phylum, in contrast to previously identified tardigrade-specific elements that are conserved in only Eutardigrada at most. This finding underlies a fundamental basis of the cellular protection against oxidative stress during anhydrobiosis and the contribution of the Golgi apparatus.

## Materials and Methods

### Tardigrade culture and UVC Exposure

In this study, we used the *Ramazzottius varieornatus* strain YOKOZUNA-1. The animals were reared in an environment established by our previous study[21] and were maintained in 90 mm petri dish plates. Approximately 600 animals were placed on 2.0% bacto-agar layered dishes and fed with *Chlorella vulgaris* mixed in Volvic water. The petri dish plates were lidded and placed in a dark incubator, set at 22 °C. The animals were transferred to new plates every 7 days.

Irradiation of the tardigrades was conducted using the UVC lamp of a drying rack (ASONE) following the protocol in our previous study[6]. Tardigrades were set on a volvic gel layered in a plastic plate, and excess water was removed. This procedure does not induce anhydrobiosis and is required to minimize UVC absorbance by the layer of water. The plates were then set for exposure to UV-C by a UVC lamp. The average dosage was 0.54 mW/cm^2^ (0.0054 kJ/m^2^), quantified by UV intensity meter (Fuso, YK-37UVSD). Water was added to the plates immediately after irradiation and set to incubate in a dark chamber set at 22°C until sampling.

### Quantification of movement in irradiated samples

10 individuals were exposed to 2.5kJ/m^2^ of UVC radiation and were placed into a single well of the 6×12 plate supplied with 2 µL of 2% agar gel. 5 replicates were prepared for UVC exposed and control individuals. A 30 second movie was taken at 0, 1, 2, 3, 6, 9, 12, 18, 24, 36, 48, 72 hours after exposure for each individual with VHX-5000 (Keyence). An individual was classified as moving if any movement was observed within this thirty-second time frame. The quantified movement was statistically compared with a non-irradiated control group by ANOVA and Tukey HSD in R. Conditions with FDR < 0.05 were classified as significant changes.

### RNA extraction and Library Preparation

We conducted RNA-Seq on two time-courses with three biological replicates, the first 0-12 hours post exposure (1h, 2h, 3h, 4h, 5h, 6h, 9h, 12h), designated “Short”, and the second 0-72 hours post exposure (0h, 12h, 24h, 36h, 48h, 72h), designated “Long”. Non-exposed individuals were sampled at 0 hours without UVC exposure. With each sampling, tardigrades were immersed in 100µL of Trizol reagent in a PCR tube (30 animals per sample) and preserved at -80°C. Biomasher II (Nippi. Inc.) was used for the homogenization of tardigrades and 200 µL of TRIzol reagent (ThermoFisher Scientific) was added to the preservation PCR tube for total RNA extraction with Direct-zol (Funakoshi). Total RNA was submitted to mRNA Isolation and RNA fragmentation with NEBNext Oligo d(T)25 beads (NEW ENGLAND BioLabs) and cDNA synthesis, adapter ligation and PCR enrichment with NEBNext Ultra RNA Library Prep Kit (NEW ENGLAND BioLabs). RNA and cDNA concentrations were determined using Qubit RNA/DNA (ThermoFisher Scientific), and electrophoresis of the synthesized library was conducted with TapeStation D1000 tape (Agilent). After pooling all of the samples (70 ng of cDNA per sample), the library was size-selected for fragments between 300-1000bp with an E-Gel EX 1% (ThermoFisher Scientific) and Nucleospin Gel and PCR Clean-up (Takara). The sequencing library was sequenced with the Illumina Next-Seq 500 instrument (Illumina) with 75 bp single end settings.

### Informatics analysis

Prior to informatics analysis, preparation of sequenced reads was conducted. After de-multiplexing sequenced raw files with bcl2fastq (Illumina), quality was checked with FastQC[36]. We then used Kallisto[37] (v0.44.0, -boot 1000, --bias) to quantify gene expression, using the *R. varieornatus* genome[13, 14]. To determine differentially expressed genes (DEGs), we mapped the RNA-Seq reads to the coding sequences with BWA MEM (v0.7.12-r1039) and conducted statistical test with DESeq2 (v1.22.2)[38, 39]. Genes with FDR below 0.05 and fold change over 2 were determined as DEGs. The expression profiles were validated using Spearman correlation in between samples and clustering. DEGs of slow-dried *R. varieornatus* anhydrobiotic samples were obtained from our previous study[13], and differentially expressed genes in both conditions were obtained.

For the identification of g12777 and g241 orthologs, we first prepared published transcriptomes and our in-house tardigrade genome database (Arakawa, Personal communication). We obtained transcriptome assemblies of *Richterius coronifer* and *Echiniscoides sigismundi* from previous studies[20]. Additionally, we re-assembled each transcriptome using Bridger[40] (v2014-12-01, *Richterius coronifer* : SRR7340056, *Echiniscoides sigismundi* : SRR7309271, default parameters). Each tardigrade genome/transcriptome was submitted to BUSCO genome completeness validation against the eukaryote lineage[41]. The gene model obtained during this assessment was used with autoAugPred.pl script in the Augustus (v3.3.3) package to predict genes in each genome[42]. The amino acid sequences of each genome were submitted to a TBLASTX or BLASTP search (v2.2.2)[43] of g12777.t1 and g241.t1 coding sequences and amino acid sequences to obtain orthologs in each species. Additionally, the g241 amino acid sequence was submitted to a Diamond BLASTP (v0.9.24.125) search against NCBI NR and NCBI Bacteria RefSeq to obtain bacterial orthologs[44, 45]. Furthermore, we submitted predicted proteome sequences for each tardigrade species and additional metazoan species used in our previous study[13] to OrthoFinder (v2.3.7, --mcl-I 2.0) to validate our BLASTP search results[46]. For g12777.t1 orthologs, amino acid sequences were subjected to MAFFT (v7.271, --auto) multiple sequence alignment and phylogenetic tree construction by FastTree (v2.1.8 SSE3, --b1000, -gamma)[47, 48], which was visualized in iTOL[49]. Orthologs were also submitted to MEME analysis (v5.1.0) for a novel motif search[50]. G-language Genome Analysis Environment (v1.9.1) was used for data manipulation[51, 52].

### Transmission electron microscope observation

Anhydrobiosis was induced by placing *R. varieornatus* in 37% RH and 0% RH for 24 hours each following our previous study[13]. Sample preparation and TEM imaging was conducted at Tokai Electronic Microscope Analysis. Samples were sandwiched between copper disks and rapidly frozen in liquid propane at -175°C. Frozen samples were freeze-substituted with 2% glutaraldehyde, 1% tannic acid in ethanol and 2% distilled water at -80°C for 2 days, -20°C for 3h, and 4°C for 4h. For dehydration, samples were incubated in ethanol for 30 minutes three times, and overnight at room temperature. These samples were infiltrated with propylene oxide (PO) twice for 30 minutes each and were put into a 70:30 mixture of PO and resin (Quetol-812; Nisshin EM Co., Tokyo, Japan) for 1h. PO was volatilized by keeping the tube cap open overnight. These samples were further transferred to fresh 100% resin and polymerized at 60°C for 48 hours before being ultra-thin sectioned at 70nm with a diamond knife using a ultramicrotome (ULTRACUT, UCT, Leica), after which sections were placed on copper grids. Sections were stained with 2 % uranyl acetate at room-temperature for 15 minutes and rinsed with distilled water. The sections were further secondary-stained with Lead stain solution (Sigma-Aldrich Co.) at room temperature for 3 minutes. The grids were observed under a transmission electron microscope (JEM-400Plusl JEOL Ltd., Tokyo, Japan) at an acceleration voltage of 100kV. Digital images (3296 x 2472 pixels) were taken with a CCD camera (EM-14830RUBY2; JEOL, Ltd., Tokyo, Japan). The images were analyzed for mitochondrial size using ImageJ and area and circularity values were tested with ANOVA and Tukey HSD in R.

### Sample preparation of recombinant proteins

For biochemical and biophysical characterizations of the g12777 protein with deletions of the N-terminal signal sequence and putative intrinsically disordered region, the gene encoding residues Gly63–Leu231 were cloned into the NdeI and EcoRI sites of the pET-28b vector (Novagen - Merck Millipore). *Escherichia coli* BL21-CodonPlus(DE3) harboring the plasmids were cultured in LB medium containing 15 µg/ml kanamycin and subsequently harvested after induction with 0.5 mM isopropyl β-D-thiogalactoside for 4 h at 37 °C. The harvested cells were resuspended with a buffer A [20 mM Tris-HCl (pH 8.0), 150 mM NaCl, and 1 mM EDTA] and lysed by sonication. The insoluble inclusion bodies were extensively washed with buffer A in the presence and absence of 2% Triton X-100, and then solubilized with 6 M guanidinium chloride, 50 mM Tris-HCl (pH 8.0), and 0.5 mM dithiothreitol. The solubilized proteins were refolded by dilution (0.05 mg/mL) in 50 mM Tris-HCl (pH 8.0), 400 mM L-arginine, 5 mM reduced glutathione, 0.5 mM oxidized glutathione, and 2 mM CaCl_2_ at 4°C for 12 h. The refolded protein was concentrated and then dialyzed against 20 mM Tris-HCl (pH 8.0), 150 mM NaCl, and 2 mM CaCl_2_. Subsequently, the N-terminal His6-tag peptide was removed by thrombin digestion. The nontagged proteins were incubated at 4°C for 30 min in the presence of 10 mM EDTA and 0.1 mM AEBSF [4-(2-Aminoethyl)benzenesulfonyl fluoride hydrochloride], and further purified with a HiLoad Superdex 75 pg (GE Healthcare) equilibrated with buffer A.

### Crystallization, X-ray data collection and structure determination

The catalytic domain of g12777 protein (10 mg/mL) was dissolved in 20 mM Tris-HCl (pH 8.0), 150 mM NaCl and 2 mM CaCl_2_. The crystals of Zn^2+^-bound g12777 protein complex were obtained in a precipitant containing 10% PEG3350, 100 mM imidazole-HCl buffer (pH 7.5), 300 mM zinc acetate, and 50 mM sodium fluoride upon incubation at 20 °C for 3 days. In contrast, the crystals of Mn^2+^-bound complex were obtained by a soaking with a buffer containing 11% PEG3350, 100 mM imidazole-HCl buffer (pH 7.5), 50 mM manganese chloride, and 50 mM sodium fluoride for 5 h using the Zn^2+^-bound crystals. The crystals were cryoprotected with the crystallization or soaking buffer supplemented with 20% glycerol. Th crystals of Zn^2+^- and Mn^2+^-bound g12777 protein complexes belonged to space groups *P*1 and *P*2_1_ with two g12777 protein molecules per asymmetric unit and diffracted up to resolutions of 1.60 Å and 2.30 Å, respectively. Diffraction data were scaled and integrated using XDS (GPLv2)[53] and AIMLESS (ver. 1.12.1)[54].

The crystal structure of the catalytic domain of the g12777 protein crystallized in the presence of excess amount of Zn^2+^ ion (300 mM) was solved by the single-wavelength anomalous dispersion (SAD) method using 1.1200 Å wavelength (Aichi-SR BL2S1, Japan) with the program Autosol in the Phenix suite (ver. 1.11.1-2575) [55]. As for the Mn^2+^-bound g12777 protein complex, the crystal structure was solved by the molecular replacement method using the Zn^2+^-bound structure as a search model. The diffraction data set was collected at Osaka University using BL44XU beamline at SPring-8 (Japan). Automated model building and manual model fitting to the electron density maps were performed using ARP/wARP (ver. 8.0)[56] and COOT (ver. 0.9)[57], respectively. REFMAC5 (ver. 5.8.0258)[58] was used to refine the crystal structure, and the stereochemical quality of the final models were validated using MolProbity (ver. 4.2)[59], showing that no amino acids were located in the disallowed regions of the Ramachandran plot. The final model of the Zn^2+^-bound g12777 protein complexes had *R*_work_ of 19.2 and *R*_free_ of 24.5%, whereas that of Mn^2+^-bound form had *R*_work_ of 26.2 and *R*_free_ of 32.5% (**Supplementary Table S2**). The molecular graphics were prepared using PyMOL (ver. 2.4.0a0, https://pymol.org/2/). These structures were compared with known protein structures using the DALI server [60]

### Measurement of enzymatic activity of the g12777 protein

The catalase/peroxidase activities of the g12777 protein were measured using Cayman Catalase Assay Kit (Cayman Chemical, USA) following the manufacturer’s protocol. For the enzymatic assay, 20 ng of the wild-type g12777 protein catalytic domain was incubated in the presence and absence of 2 mM ZnCl_2_ and MnCl_2_ for 30 min at 25 °C. Bovine liver catalase (Sigma Aldrich) was used as a positive control. Catalase/peroxidase activity were tested with ANOVA and Tukey HSD in R.

### Isothermal titration calorimetry

Purified catalytic domain of the g12777 protein dissolved in 20 mM Bis-Tris-HCl (pH 6.5) containing 150 mM NaCl was used for isothermal titration calorimetry (ITC). In this experiment, a syringe containing 1 mM ZnCl_2_, MnCl_2_ or CaCl_2_was titrated into a sample cell containing 0.1 mM catalytic domain of the g12777 protein using an iTC200 calorimeter (GE Healthcare).

### Expression of GFP-tagged g12777 in HEK293

For expression of GFP-fused g12777 proteins, we inserted the coding sequence between the *Sal*I and *BamH*I restriction sites of the pAcGFP1-N1 plasmid (Takara). The total RNA was extracted from *R. varieornatus* with Direct-zol Ultra RNA (Funakoshi) and was reverse transcribed with PrimeScript Reverse Transcriptase (Takara). g12777 mRNA was selected by PCR using Tks Gflex DNA Polymerase (Takara) with the following primers:

g12777-F: 5’-ATTGTCGACATGGCATTATCTTTGTGGATGACTG-3’,

g12777-R: 5’-TAAGGATCCTTTAGGGAAGAAGGTGCCCGACAG-3’.

For construction of Δ62aa construct, the forward primer was changed to 5’-ATTGTCGACGGCCGCTTTGCCGATTTCTTCAGAA-3’. The corresponding g12777-D2A (D92A, D98A, D161A, D163A mutations, AMNP-D2A) coding sequences were adapted for human codon usage and synthesized at Eurofin Genomics. g12777-D2A was inserted into the pAcGFP1-N1 plasmid (Takara) between and *Sal*I-*BamH*I. Constructs were transfected into HEK293 cells with the X-tremeGENE 9 reagent (Sigma-Aldrich) and submitted to selection with G418 (Sigma-Aldrich) at 400 µg/mL for more than two weeks, and further passaged at 100 µg/mL. HEK293 cell lines were cultured with MEM medium (Sigma-Aldrich) supplied with FBS (Funakoshi), NEAA (ThermoFisher Scientific) and Antibiotic-Antimycotic Mixed Stock Solution (Nacalai) and were passaged every 3-4 days with TryPLE (ThermoFisher Scientific). For microscopy observations, cells were fixed with 4% Paraformaldehyde (Wako), and co-stained with 300 nM DAPI (ThermoFisher Scientific) and 20 µL of CellLight Golgi RFP, BacMan 2.0 (ThermoFisher Scientific) to visualize the nucleus and Golgi apparatus, respectively. Fluorescent signals were observed under the SZ-5000 (Keyence) at 40x or 100x magnification.

Additionally, the localization of six high expressed AMNP orthologs were validated using similar methods. cDNA was obtained from *R. varieornatus* with RNeasy Mini Kit (Qiagen) and PrimeScript II 1st strand cDNA Synthesis Kit (Takara). The coding sequences of the corresponding genes were amplified by PrimeSTAR Max DNA Polymerase (Takara) using the following primers: g243-F: 5’-ATTCTGCAGTCGACGGTATGCAGGATGTTTCCGA-3’,

g243-R: 5’-catgaccggtggatcCTATTTGGTGAACCATCCG-3’,

g244-F: 5’-ATTCTGCAGTCGACGATGGCCAAGGCGGCAATC-3’,

g244-R: 5’-catgaccggtggatcTCATCTAGAGAAAAGTCCGC-3’,

g245-F: 5’-ATTCTGCAGTCGACGGTATGAACTTTCTCTGCTGG-3’,

g245-R: 5’-catgaccggtggatcTCAGGAGAACATGCCCGA-3’,

g246-F: 5’-ATTCTGCAGTCGACGATGATGCAGCTGACAATCTT-3’,

g246-R: 5’-catgaccggtggatcTTATGAGAACATGCCGTCG-3’,

g779-F: 5’-ATTCTGCAGTCGACGGTATGGATCTGGACAGGG-3’,

g779-R: 5’-catgaccggtggatcTCATTTTCCAATAAAGGGAATG-3’,

g2856-F: 5’-ATTCTGCAGTCGACGGTATGTGGGGAATACTGTG-3’,

g2856-R: 5’-catgaccggtggatcTCATGCCATAGCGTGGCG-3’,

g12777-F: 5’-ATTCTGCAGTCGACGGTATGGCATTATCTTTGTGG-3’,

g12777-R: 5’-catgaccggtggatcTTATAGGGAAGAAGGTGCC-3’,

These coding sequences were inserted into the pAcGFP1-N1 plasmid by In-Fusion HD Cloning Kit. Constructs were transfected into HEK293 cells with the X-tremeGENE 9 reagent (Sigma-Aldrich) in optiMEM solution for 4 hours. After incubation for 24-48 hours in MEM medium, cells were co-stained with Hoechst33342 (ThermoFisher Scientific) and CellLight Golgi-RFP (ThermoFisher Scientific) and observed under the Leica SP8 microscope with x100 magnification.

### Validation of cellular tolerance against oxidative stress

To evaluate cellular tolerance of AMNP transfected cells, we conducted MTT analysis of H_2_O_2_ exposed cell lines expressing only GFP, g12777, g12777-Δ62aa, and g12777-D2A (3 technical replicates). In brief, approximately 10,000 cells were plated into 96 well plates and were incubated at 37°C 5% CO_2_ for 24 hours. Cells were exposed to 0-1.0 mM H_2_O_2_ (Wako) for 30 minutes (3-5 replicates). The culture medium was removed to stop exposure, and 100 µL MEM medium was added to the wells and further incubated for 24 hours. The culture medium was replaced with 100 µL fresh culture medium supplied with 10 µL of 5 µg/µL Thiazolyl Blue Tetrazolium Bromide (Sigma-Aldrich) and incubated for 2 hours for formazan formation. The culture medium was removed, and 100 µL DMSO (Wako) was added to melt the formazan. The 96 well plate was measured with 570 nm and 670 nm using SPECTRAmax PLus384 (Molecular Devices). Three technical replicates were measured. Absorbance values were calculated by Ab570 – Ab670 – Ab-BLANK. Absorbance values were statistically tested with ANOVA and Tukey HSD in the R program.

For MACS flow cytometry, cells were cultured and exposed to H_2_O_2_ similar to the MTT assay protocol. After 24-hour incubation after exposure, the MEM medium (100 µL) was moved to a clean well and each well was washed with 100 µL PBS (-). All of the PBS was moved to the corresponding well. 50 µL TryPLE was added for trypsinization and incubated at 37°C in the CO_2_ incubator for 4 minutes. The MEM culture and PBS mixture were moved to their original wells and mixed thoroughly to release the cells. The 96 well plate was centrifuged at 2,000 rpm (750g) for 6 minutes at 4°C, and the supernatant was removed. Each well was washed with 1x Annexin binding buffer (ThermoFisher Scientific), and was centrifuged at 2,000 rpm (750g) for 6 minutes at 4°C. The buffer was removed, and each well was supplied with 100 µL of Annexin binding buffer supplied with 0.1 µL SYTOX™ Blue Nucleic Acid Stain (ThermoFisher Scientific) and 2 µL Annexin V Alexa Fluor 657 (ThermoFisher Scientific). The cells were resuspended with pipetting and set to stain for 10 minutes on ice. The MACS Quant10 instrument (Miltenyi Biotec) was set for measurement of DAPI, Alexa 657 and GFP. Three technical replicates were measured. Approximately 100-1500 cells were measured for each cell line. The ratio of healthy cells (Anninx-V (-) SYTOX blue (-) cells) were compared with ANOVA and Tukey HSD in the R program. Adjusted p-values (FDR) below 0.05 were designated as significant differences.

## Acknowledgments

We thank Nozomi Abe and Naoko Ishii for tardigrade sample preparation. We also thank Yuki Takai, Dr. Akio Kanai, Fumie Nakasuka, Dr. Shojiro Kitajima and Dr. Sho Tabata (Keio University IAB) for their advice on the experiments. We acknowledge Dr. Maho Yagi-Utsumi (ExCELLS) for her useful discussion. We thank Kumiko Hattori (Nagoya City University) for their help in the preparation of recombinant proteins. We also thank Dr. James Fleming (Keio University IAB) for proofreading the manuscript. The diffraction data set were collected at Nagoya University using the BL2S1 beamline at Aichi Synchrotron Radiation Center (Japan) and Osaka University using BL44XU beamline at SPring-8 (Japan). We thank the beamline staff for providing the data collection facilities and support. We acknowledge the assistance of the Research Equipment Sharing Center at Nagoya City University for ITC measurement. This work is supported by KAKENHI Grant-in-Aid for Scientific Research (B) and Grant-in-Aid for JSPS Fellows from the Japan Society for the Promotion of Science (JSPS, grant no. JP18J21155, 17H03620), Joint Research by Exploratory Research Center on Life and Living Systems (ExCELLS program No. 19-208 and 19-501) and partly by research funds from the Yamagata Prefectural Government and Tsuruoka City, Japan. *C. vulgaris* used to feed the tardigrades was provided courtesy of Cholorela Industry.

## Competing interest declaration

All authors do not have any competing interests.

## Data availability

The coordinates and structural factors of the crystal structure of the catalytic domain of g12777 protein complexed with Mn^2+^ or Zn^2+^ ions have been deposited in the Protein Data Bank under accession numbers 7DBT and 7DBU, respectively. The RNA-Seq data obtained were deposited to NCBI GEO under the accession ID GSE152753.

---

**S1 Table: Statistics of RNA-Seq data obtained in this study**. Statistics of each RNA-Seq data sequenced in this study. Mapping ratio for BWA mapping are shown.

**S2 Fig.**
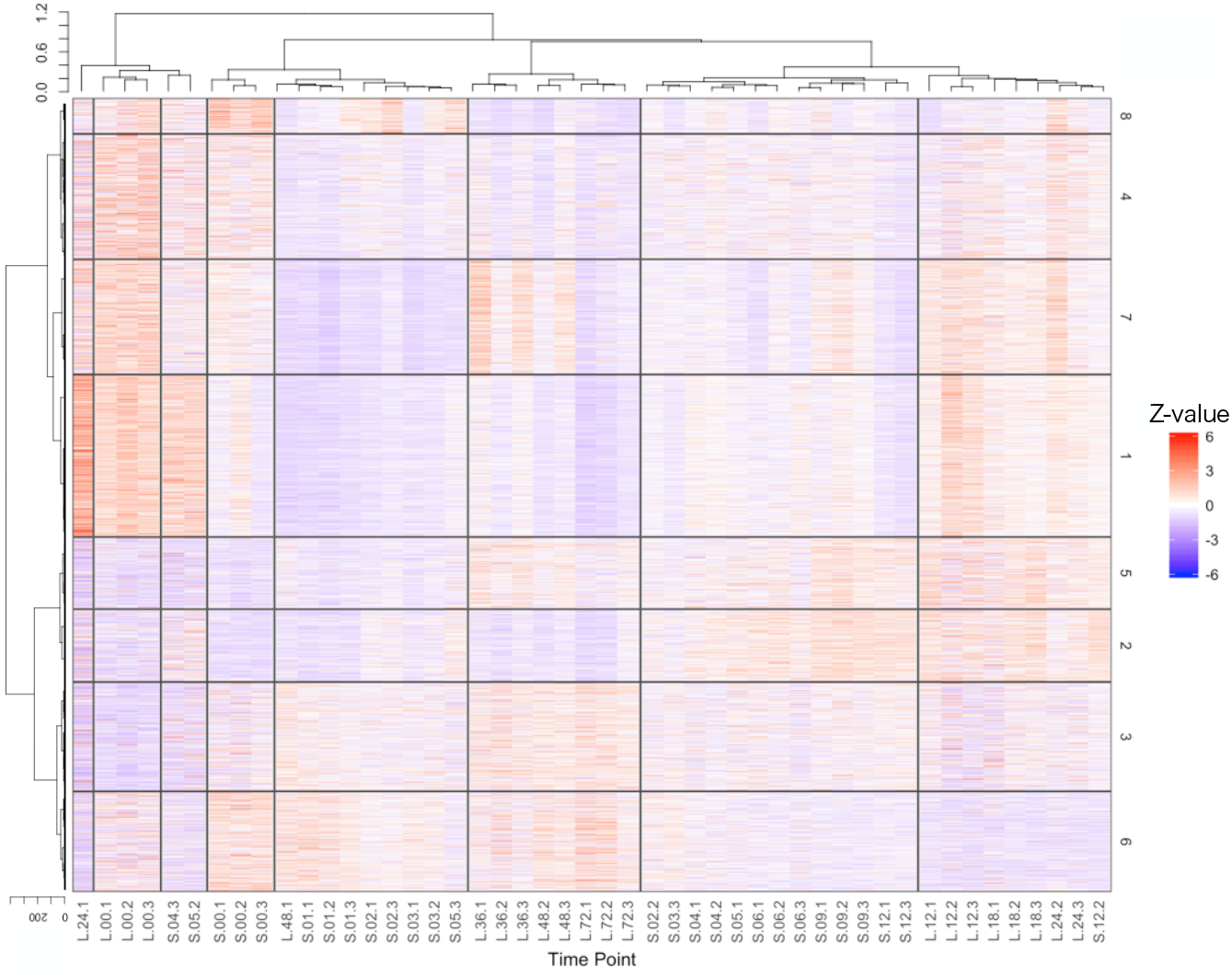
Expression profiles of UVC exposed *R. varieornatus*. Gene expression profiles of *R. varieornatus* specimens exposed to 2.5kJ/m^2^ UVC shown as a heatmap. TPM values were Z-scaled to between samples. Expression and sample profiles were clustered with by Ward method based on Spearman correlation.

**S3 Table. Highly expressed or regulated genes in Short time-course**. Genes with TPM values > 1000 and fold change > 4 are highlighted in purple and orange, respectively.

**S4 Table. Highly expressed or regulated genes in Long time-course**. Genes with TPM values > 1000 and fold change > 4 are highlighted in purple and orange, respectively.

**S5 Text. Abnormal mitochondrial structure in recovering specimens**. We found mitochondria-related genes, in particular, the mitochondrial chaperone BCS1 and the mitophagy related gene Sequestosome, to have a high fold change. We have found similar inductions during *H. exemplaris* anhydrobiosis, suggesting the existence of mitochondria stresses during desiccation or UVC exposure-response. Oxidative stress caused during desiccation (or UVC exposure) would also induce extensive stress to the mitochondria. We conducted TEM observations of *R. varieornatus* specimens recovering from desiccation. We observed abnormal mitochondrial morphology in specimens 3 hours after rehydration, not in 30 minutes (**S6 Fig**). This suggests that mitochondrial stress may be occurring during the initial 3 hours, resulting in morphological changes at 3 hours after rehydration. Induction of Sequestosome may be related to mitophagy of damaged mitochondria. A previous study using *H. exemplaris* specimens also has observed mitochondrial morphology, however, they have seen an increase in mitochondrial size, not a decrease as we have observed [61]. These inconsistencies may reflect the differences in mitochondrial damage that occurred during anhydrobiosis in the two species.

**S6 Fig.**
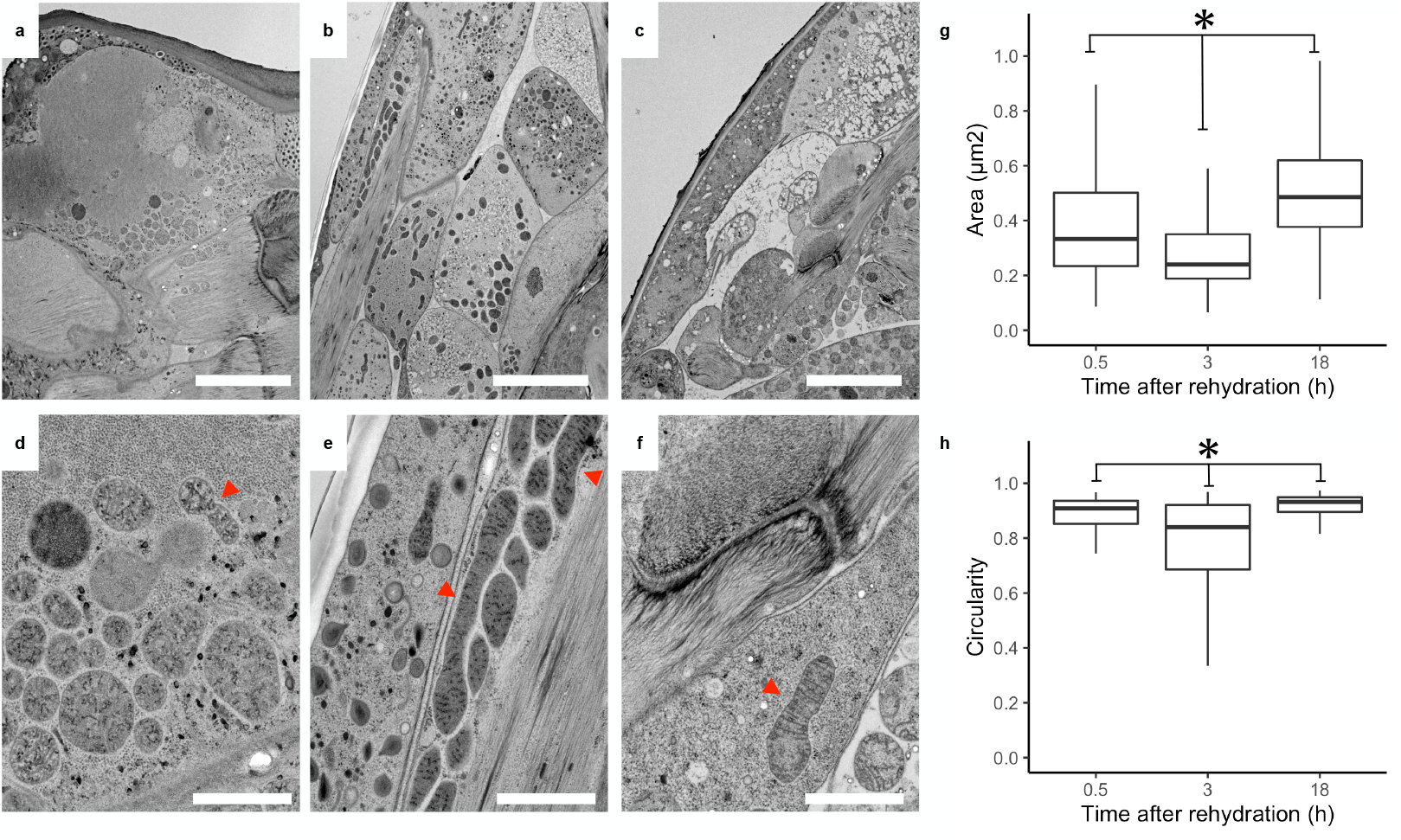
Abnormal mitochondrial structures in anhydrobiosis recovering *R. varieornatus*. Transmission electron microscopy images of *R. varieornatus* specimens during recovery from anhydrobiosis. [ad]: 0.5h [be]: 3h, [cf]: 18h post recovery. [abc]: 42,00x magnification of mitochondria. Scale bar 5µm. [def]: 21,000 magnification of mitochondria. Abnormal mitochondria structures can be observed (Red arrows). Scale bar 1µm. [gh] Decrease in area [g] and circularity [h] values in individuals recovering from anhydrobiosis. Mitochondria structures were outlined in ImageJ (0.5h: 159, 3h: 355, 18h: 154 structures) and was statistically tested (ANOVA + Tukey HSD, FDR<0.05).

**S7 Table. List of differentially expressed genes**. List of genes differentially expressed in both UVC response and slow-dry desiccation. Known anhydrobiosis related genes are highlighted in orange.

**S8 Table. Enrichment analysis of DEGs**. Enrichment analysis of KEGG pathway, Pfam-A and Gene ontology term of differentially expressed genes. Terms colored in red are terms suggested to be related to anhydrobiosis in previous studies

**S9 Text. Details on identification of novel stress-responsive gene families**. To validate the conservation within Tardigrada, we submitted gene sequences to a BLAST search against our in-house and publicly accessible genomes/transcriptomes. We first predicted approximately 10-90 thousand genes in various tardigrade lineages and then identified orthologs by BLASTP search or OrthoFinder clustering (**S12 Table, S13 Table, S14 Table**). Most of the linages showed conservation of g2856 (excluding Halechiniscidae family in Arthrotardigrada), indicating that this gene family is highly conserved in the phylum (**S12 Table, S13 Table**). On the contrary, we found that g241 was lost in the Apochela, Echiniscoididae and Arthrotardigrada linages (**S12 Table, S14 Table**). Additionally, initial TBLASTX searches against the publicly accessible *Echiniscoides sigismundi* and *Richterius coronifer* transcriptome assemblies indicated the loss of g2856 orthologs both species. We determined several raw RNA-Seq reads that showed homology against g2856 coding sequences, which implied the existence of g2856 orthologs. Therefore, we re-assembled both the *E. sigismundi* and *R. coronifer* transcriptome using previously sequenced RNA-Seq data with Bridger. BLASTX searches against this re-assembly found approximately 15 g2856 orthologs but no g241 orthologs in *E. sigismundi*. On the other hand, we detected 23 g241 and 64 g2856 orthologs in *R. coronifer*. The lack of g241 or g2856 orthologs in previous transcriptome assemblies may have occurred during the filtering stage. Our in-house transcriptome sequencing data supported the existence of both gene families in *R. coronifer* (56 g2856 and 17 g241 orthologs) and 12 g2856 orthologs in *E. sigismundi*. Together, these results support that the g241 is lost in the Echiniscoididae, Arthropoda, and Apochela lineages. Three of the g2856 orthologs (including g2856) were found to be mispredicted in our gene set (**S15 Table**).

**S10 Table. Expression of g241**.**t1**. TPM values of g241 orthologs in *R. varieornatus* UVC response.

**S11 Table. Expression of AMNP**. TPM values of g12777 orthologs in *R. varieornatus* UVC response.

**S12 Table. AMNP and g241**.**t1 orthologs within Tardigrada identified by BLASTP**. Coding sequences of g241.t1 and g12777.t1 were submitted to BLASTP search against predicted proteome sequences of indicated species in our in-house genome database and additional transcriptome assemblies.

**S13 Table. AMNP copy numbers within Tardigrada identified by Orthofinder**. Number of genes classified as g12777 orthologs by Orthofinder.

**S14 Table. g241 copy numbers within Tardigrada identified by Orthofinder**. Number of genes classified as g241 orthologs by Orthofinder.

**S15 Table. Refinement of AMNP gene regions**. Mispredicted AMNP orthologs in *R. varieornatus* genome.

**S16 Fig.**
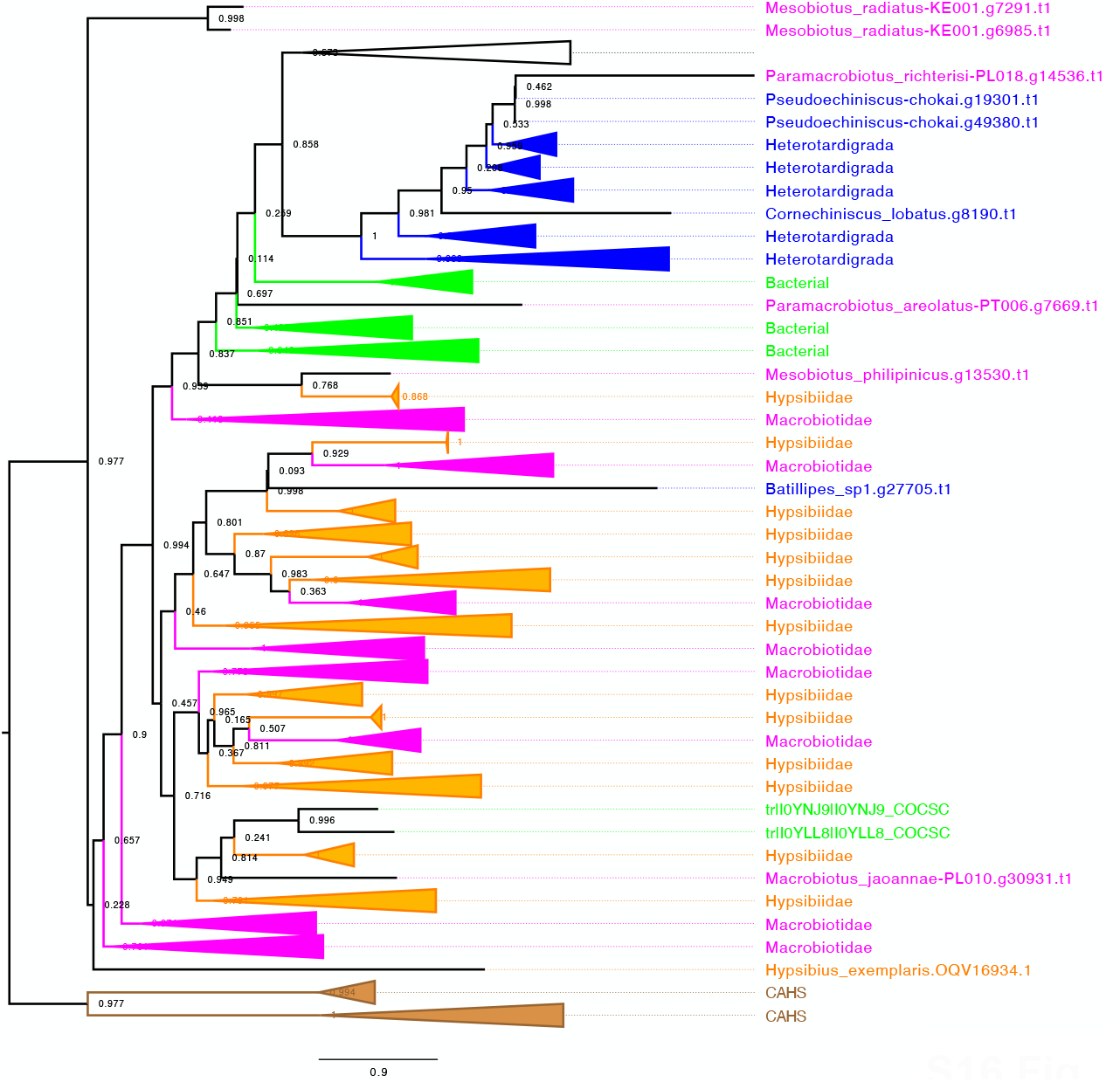
Phylogenetic analysis g241 orthologs within Tardigrada. Phylogenetic tree of tardigrade g241 orthologs and bacterial orthologs. Amino acid sequences were aligned with Mafft and the phylogenetic tree was constructed with FastTree with 1000 bootstraps (-boot, -gamma). CAHS genes were used as an out-group.

**S17 Text. Categorization of g241 orthologs**. The g241 orthologs in *H. exemplaris* were identified as a high-confidence horizontally transferred gene candidate in our previous study (**S18 Table**). BLASTP searches against TrEMBL and NCBI nr databases indicated that the majority of homologs originates from Gammaproteobacteria species (**S19 Table, S20 Table**). This suggests that this gene family may have been integrated into the tardigrade genome before the divergence of these Eutardigrada and Heterotardigrada. We also have found several orthologs in Ciliophora, Chlorophyta, and Acanthamoeba, however, the conservation patterns suggest that these genes may also be results of horizontal transfer into these organisms. Several of these bacterial orthologs were found to be fused to the C-terminal end of haem peroxidases. These regions do not have any functional domains and are not predicted to be a disordered region, suggesting that this gene itself may not have anti-oxidative stress

**S18 Table. HGT statistics of g241 orthologs from previous studies**. HGT statistics calculated in our previous study[13] for g241 orthologs.

**S19 Table. g241**.**t1 orthologs identified from NCBI nr database**. List of orthologs identified by Diamond BLASTP search (E-value < 1E-5) against NCBI nr database. Tardigrade, non-tardigrade eukaryotic, and bacterial hits were each colored in gray, yellow, and green. *H. exemplaris* orthologs colored in red had gene annotations, but validation by RNA-Seq data mapping indicated that these genes may be mis-predicted.

**S20 Table. g241**.**t1 orthologs identified from RefSeq database**. List of orthologs identified by Diamond BLASTP search (E-value < 1E-5) against NCBI Bacteria RefSeq complete genome sequences. Genes before and after g241 orthologs are shown.

**S21 Table. Syntenic information of g2856 and g241 orthologs in *R. varieornatus***. Annotation of 5 genes prior and after g2856 orthologs in *R. varieornatus*. g2856 and g241 orthologs are highlighted in red.

**S22 Fig.**
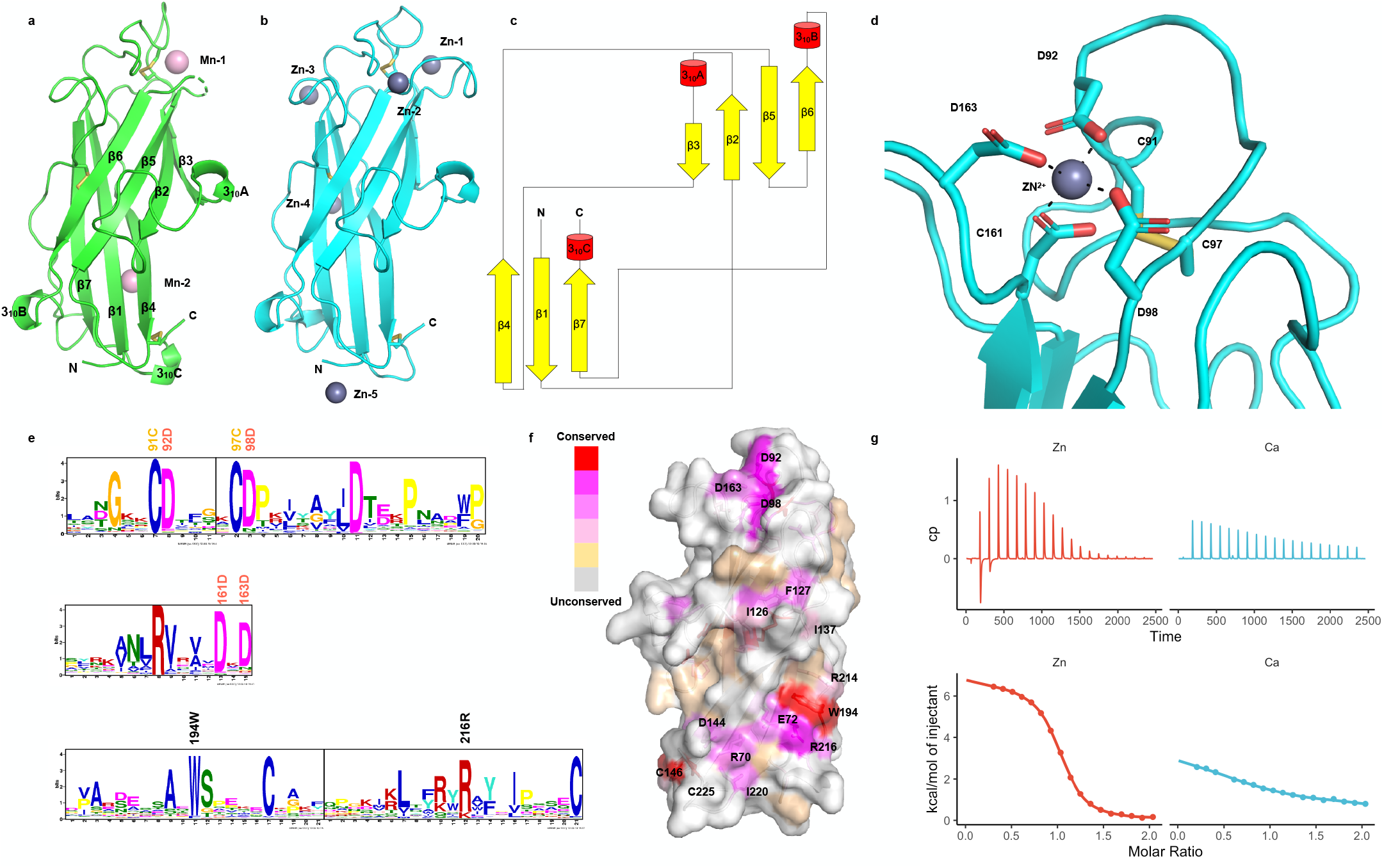
Metal-binding property and structural detail of g12777 globular domain. [a,b] The crystal structures of catalytic domain of g12777 complexed with Mn^2+^ [a] and Zn^2+^ [b]. All crystallographically observed metal ion binding sites are indicated. [c] Topology diagram of g12777 catalytic domain. [d] Close-up view of the Zn^2+^-binding site. The residues comprising binding site 1 (D92, D98, D161, and D163) and the disulfide bond (C91 and C97) are indicated. [e] Conservation of residues. All g12777 orthologs were submitted to MEME search. [f] Conserved resides of g12777. The conserved resides are colored according to the bit scores obtained from MEME analysis. [g] Ca^2+^ and Zn^2+^ ion binding affinity measured by isothermal titration calorimetry. The upper two panels indicate the raw data, while the bottom two panel represent the integrated heat values corrected for the heat of dilution and fit to a one-site binding model (solid line); red: Ca^2+^, cyan: Zn^2+^.

**S23 Table: Data collection and refinement statistics for g12777**. Crystal parameters and refinement statistics for Mn^2+^- and Zn^2+^-bound g12777 protein catalytic domain are summarized. *One crystal for each structure was used for diffraction data collection.

**S24 Text: Details on the crystal structure of g12777 protein**. In the Zn^2+^-bound crystal structure, five Zn^2+^ ions (Zn-1-5) originated from the crystallization buffer containing 300 mM zinc acetate were observed, while Ca^2+^ originated from the protein buffer (2 mM calcium chloride) was not. We prepared Mn^2+^-bound g12777 crystals by soaking method using Zn^2+^-bound crystals. Upon soaking of excess amount of Mn^2+^ (50 mM), Zn^2+^ was replaced with Mn^2+^ in Mn-1 and newly appeared in Mn-2 (**S21a Fig**). On the other hand, Zn^2+^ in Zn-2-5 disappeared in the Mn^2+^-bound crystal structure. As predicted from the bioinformatics analysis, the crystal structure of putative catalytic domain (residues Gly63–Leu231) of the g12777 protein displayed a characteristic β-sandwich fold comprising seven β-strands and three 3_10_ helices with three disulphide bridges (C91-C97, C146-C225, and C176-C200) (**Fig 2a, S21bc Fig**). In the crystal structure of Mn^2+^-bound form, a part of β5–β6 loop (residues 163–168) around the Mn^2+^-binding site was disordered, suggesting its flexible nature. Comparison of the structure of the β-sandwich domain of g12777 with known protein structures revealed that the g12777 protein structure has very weak similarities with structures of calcium-binding C2 domain involved in lipid interaction (*Z*-score = 5.1–5.5; RMSD = 2.5–3.2 Å; identity = 5–13%; PDB codes: 2JGZ, 1WFJ, and 4IHB). In addition, it has subtle similarity with a Cu/Zn-SOD monomeric circular permutant (*Z*-score = 4.1; RMSD = 3.4 Å; identity = 5%; PDB code: 5J0C).

**S25 Table. Metal ion affinity of recombinant proteins**. Affinity values for metal ions with g12777 proteins.

**S26 Fig.**
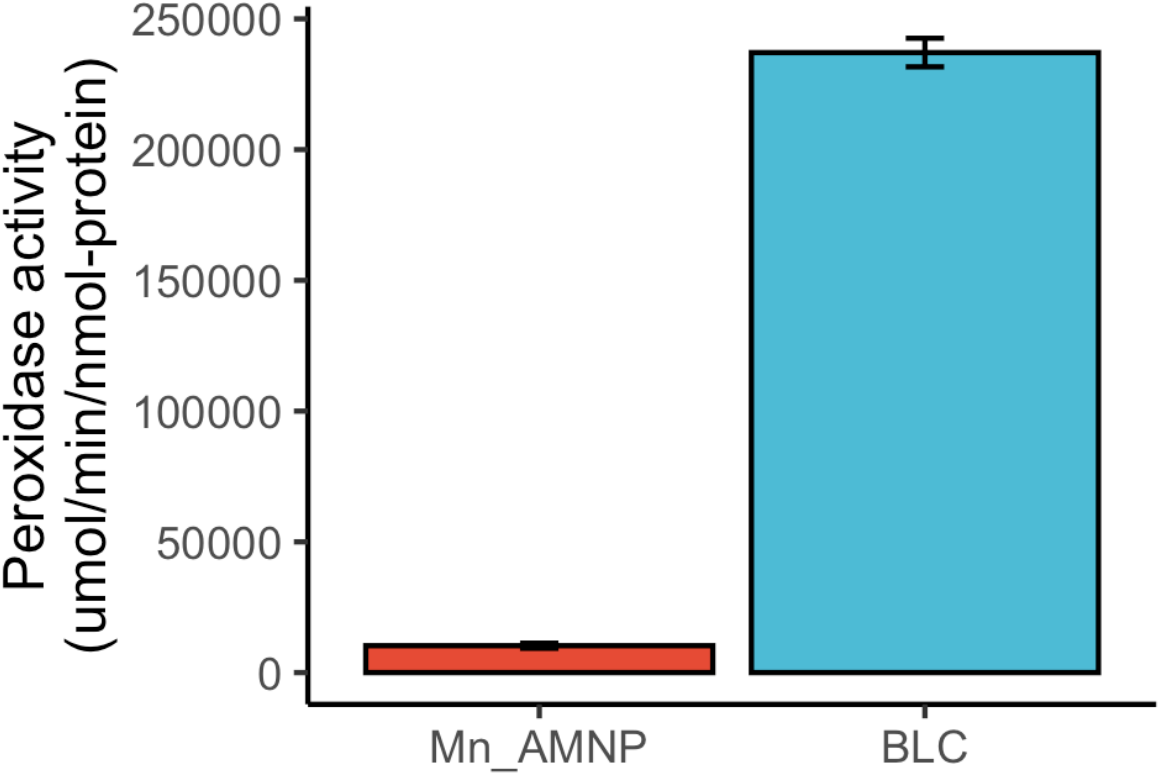
Localization of other AMNP orthologs. The localization of top six high expressed AMNP orthologs were validated using the same method. All of these orthologs showed localization to the Golgi apparatus.

**S27 Fig.**
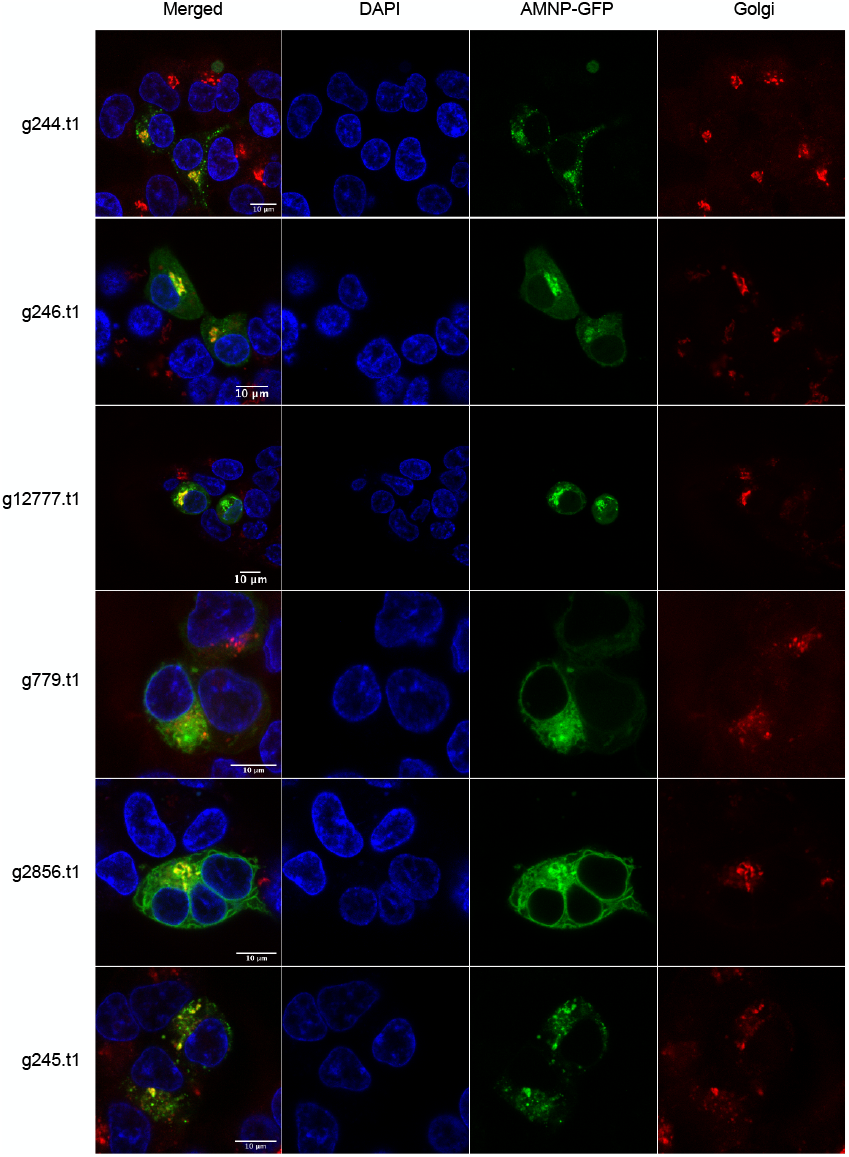
Comparison of peroxidase function with Bovine liver catalase. The peroxidase function of AMNP was compared to those of Bovine liver catalase.

## Notes

### Competing Interest Statement

The authors have declared no competing interest.

